# Generalizability of foot-placement control strategies during unperturbed and perturbed gait

**DOI:** 10.1101/2023.07.10.548298

**Authors:** Chang Liu, Francisco J. Valero-Cuevas, James M. Finley

## Abstract

Control of foot placement is an essential strategy for maintaining balance during walking. During unperturbed, steady-state walking, foot placement can be accurately described as a linear function of the body’s center of mass state at midstance. However, it is uncertain if this mapping from center of mass state to foot placement generalizes to larger perturbations that may be more likely to cause falls. These perturbations may cause balance disturbances and generate reactive control strategies not observed during unperturbed walking. Here, we used unpredictable changes in treadmill speed to assess the generalizability of foot placement mappings identified during unperturbed walking. We found that foot placement mappings generalized poorly from unperturbed to perturbed walking and differed for forward versus backward perturbations. We also used singular value decomposition of the mapping matrix to reveal that people were more sensitive to backward versus forward perturbations. Together, these results indicate that control of foot placement during losses of balance differs from the control strategies used during unperturbed walking. Better characterization of human balance control strategies could improve our understanding of why different neuromotor disorders result in heightened fall risk and inform the design of controllers for balance-assisting devices.

## 1 Introduction

Control of foot placement is an important strategy for maintaining balance during walking [1] [2] [3] [4]. Balance can be controlled via foot placement by varying the center of pressure and the magnitude of the ground reaction force to influence the body’s linear and angular momentum. For example, one way to recover from a forward loss of balance is to place the foot more anterior to the body’s extrapolated center of mass (CoM) than normal. This strategy produces a ground reaction force that has a greater posteriorly-directed component to reduce forward linear momentum while also producing a backward moment about CoM to arrest the forward rotation of the body [5]. Thus, modulating foot placement from step-to-step is an important strategy for humans to maintain balance.

Step-to-step balance corrective strategies are often characterized using a data-driven approach relating foot placement location to the body’s state at an earlier phase of the gait cycle [6] [7] [8] [9] [10] [11] [12] [13] [14]. Given an average CoM trajectory and many strides of steady walking, one can often derive a linear mapping between deviations of the CoM state from this trajectory to deviations in the next foot placement [8] [11] [12]. These mappings can explain *∼*80% of the variance in foot placement in the mediolateral direction and *∼*30 - 40% of the variance in the anteroposterior direction using the CoM state at midstance [12] [13] [15]. Though passive dynamics may lead to some degree of correlation between CoM state and foot placement [16], the high degree of variance explained, especially in the mediolateral direction, may indicate that the central nervous system uses information about the body’s state to actively control the next foot placement during unperturbed walking.

Although the observed mappings explain foot placement patterns during unperturbed gait, the extent to which these mappings generalize to perturbed walking remains to be seen. It is conceivable that linear mappings may fail to explain balance correcting responses to external perturbations and if so, this would suggest that studying unperturbed walking alone is insufficient for elucidating the strategies that people use to prevent falls. Recently, the generality of a linear mapping between deviations in CoM state and subsequent foot placement has been examined using intermittent backward perturbations [6]. In this study, approximately 30% of the variance in fore-aft foot placement was explained by a linear mapping derived from perturbed steps [6]. However, it has yet to be determined how or if this mapping differs from that inferred from steady-state, unperturbed walking or if foot placement strategies differ for backward versus forward perturbations.

The primary goal of this study was to determine whether the mapping between CoM state and foot placement derived from unperturbed walking could explain the variance in foot placement in response to forward and backward perturbations in neurotypical adults. We hypothesized that a mapping that accounted for the directional differences in response to unexpected forward versus backward disturbances would better explain the variance in foot placement than a mapping derived solely from unperturbed walking. This is because one might expect different strategies to be effective when balance disturbances are in the same versus the opposite direction of linear momentum. Additionally, we performed singular value decomposition on the foot placement mapping to provide a direct assessment of the direction along which foot placement was most sensitive to deviations in CoM state and the sensitivity of foot placement control along that direction. We expected to find differences in the derived foot placement mappings as well as the direction and sensitivity of foot placement control to deviations in CoM state between unperturbed and perturbed walking. Overall, this study may extend our understanding of how people control foot placement to maintain balance during walking and may inform the design of controllers for assistive devices to stabilize walking in response to perturbations.

## 2 Materials and methods

### 2.1 Participant characteristics

A total of 13 neurotypical adults with no musculoskeletal or gait impairments participated in this study (6F, 58 ± 29yrs, 0.75 ± 0.25 m/s). These participants were recruited as age-matched controls for a sample of post-stroke participants from a prior study [5]. All participants reported their right side as their dominant limb when asked which leg they would use to kick a ball. The study was approved by the Institutional Review Board at the University of Southern California (#HS-18-00533), and all participants provided informed consent before participating. All aspects of the study conformed to the principles described in the Declaration of Helsinki.

### 2.2 Experimental protocol

Participants walked on an instrumented, dual-belt treadmill (Fully Instrumented Treadmill, Bertec, USA) for six separate trials at their self-selected walking speed. We determined their self-selected walking speed using a two-alternative forced-choice staircase method [17] [18] [19] as described in [20]. Participants then walked on the treadmill for five minutes at their self-selected walking speed without receiving any perturbations. Then, for five subsequent trials, participants reacted to acceleration of the treadmill belts. Each trial consisted of a total of 24 perturbations with 12 on each belt. The perturbations had magnitudes of -0.5 m/s, -0.4 m/s, -0.3 m/s, 0.3 m/s, 0.5 m/s, and 0.7 m/s, where positive values indicate increases in speed relative to the participant’s self-selected walking speed, and negative values correspond to reductions in the participant’s self-selected walking speed. Each perturbation was remotely triggered by customized Matlab code and the order of these perturbations was randomized. Each perturbation was characterized by a trapezoidal speed profile in which the treadmill accelerated at the time of foot strike to the target belt speed at an acceleration of 3 m/s^2^ (or -3 m/s^2^ if the target speed was less than their walking speed), held this speed for 0.7 s, and then returned to the participant’s self-selected walking speed at an acceleration of -3 m/s^2^ (or 3 m/s^2^) [21]. The perturbations were randomly triggered to occur within a range of 15 to 25 steps after the previous perturbation to provide participants with sufficient time to reestablish their baseline walking pattern and prevent them from anticipating perturbation timing.

### 2.3 Data Acquisition

We used a ten-camera motion capture system (Qualisys AB, Gothenburg, Sweden) to record 3D marker kinematics at 100 Hz and ground reaction forces at 1000 Hz. We placed a set of 14 mm spherical markers on anatomical landmarks and marker clusters on the upper arms, forearms, thighs, shanks, and the back of heels to create a 13-segment, full-body model [22] [23]. We calibrated marker positions during a five-second standing trial and removed all joint markers after the calibration.

### 2.4 Data Processing

We post-processed the kinematic and kinetic data in Visual3D (C-Motion, Rockville, MD, USA) and Matlab 2020b (Mathworks, USA) to compute variables of interest. We lowpass filtered marker positions and ground reaction forces using 4th order Butterworth filters with cutoff frequencies of 6 Hz and 20 Hz, respectively, based on previous literature [24] [25] [26]. Foot strike was defined as the time point when the ground reaction reached 80N. We also examined the timing of perturbations relative to foot strike post-hoc to remove the perturbations that occurred more than 150ms after the foot-strike [27]. We included a median of 10 (interquartile range: 1) perturbations for each perturbation amplitude per side for each participant.

### 2.5 Models of Foot Placement

Our goal was to derive a mapping between CoM state and foot placement to characterize the step-to-step balance corrective strategies during unperturbed and perturbed walking. The CoM state during single limb stance, ***s***, was defined as in Eqn.1.

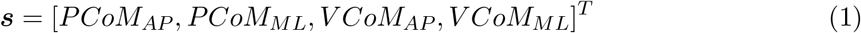

The position of the next foot placement ***q*** was defined as in Eqn.2.

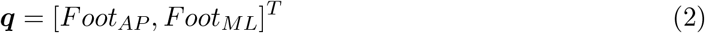

CoM state included the CoM position (PCoM) and velocity (VCoM) in the fore-aft (AP) and mediolateral (ML) direction. Both CoM state and foot placement positions were relative to the position of the current stance foot (Figure 1). We normalized position variables using the height (H) of the participant and velocity variables using 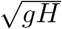 where g is the gravity constant. Each step cycle was divided into 100 time points.

**Figure 1.**
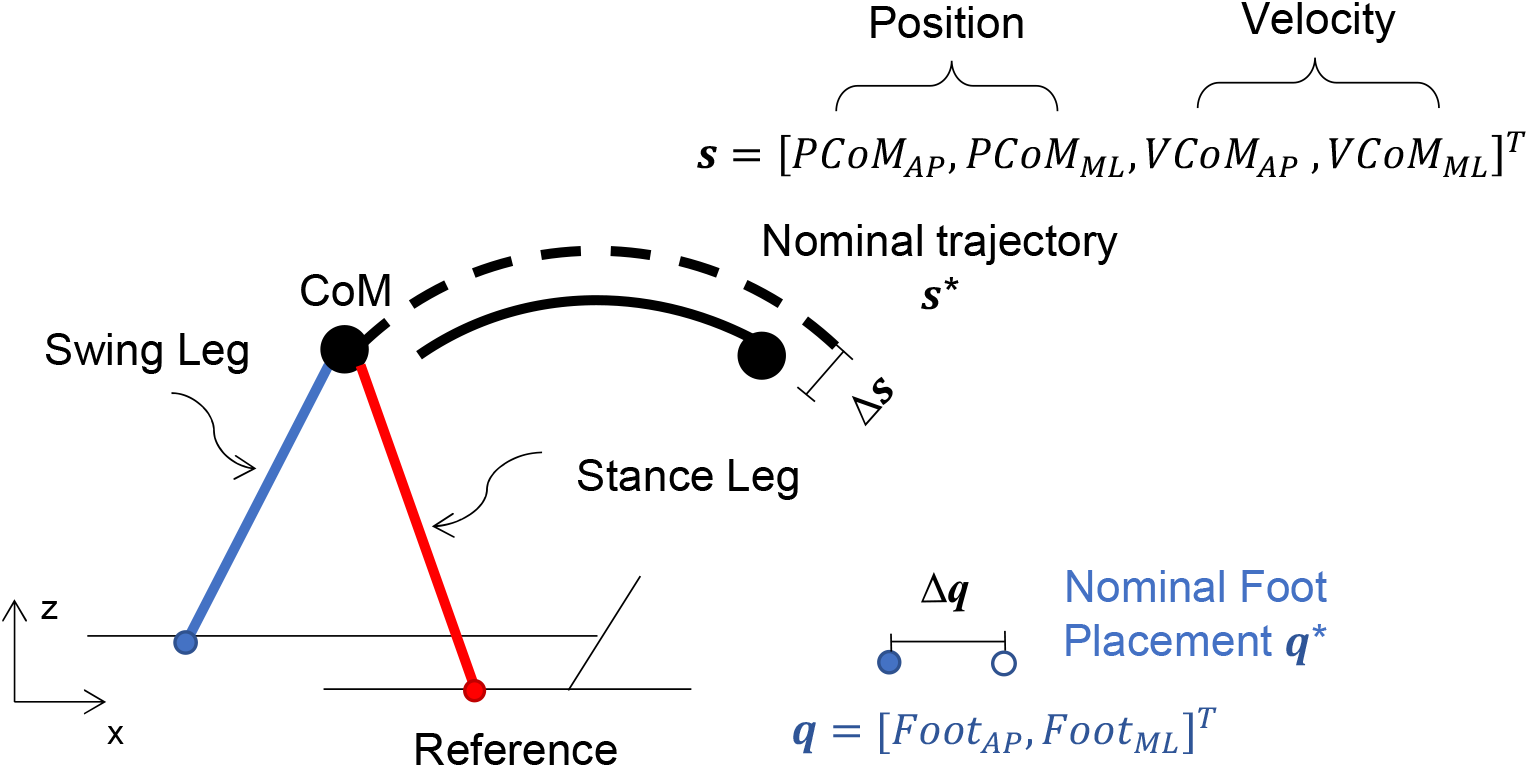
Diagram of the model describing the CoM state (*s*) and foot placement (*q*). CoM state included the CoM position (PCoM) and velocity (VCoM) in the fore-aft (AP) and mediolateral (ML) direction. Blue: swing leg, Red: stance leg. CoM position and the position of the swing foot were referenced to the stance foot. The black dashed trajectory represents the nominal (average) CoM trajectory. The black solid trajectory represents one measured trajectory. Δ***q*** and Δ***s*** represent the step-to-step fluctuation of the foot placement and CoM state. AP: anteroposterior; ML: mediolateral.

We defined the nominal trajectories of the CoM (***s***^***∗***^) and foot-strike positions (***q***^***∗***^) as the average values of these quantities during unperturbed walking. Step-to-step fluctuations about the nominal trajectory allowed us to determine the relationship between deviations in foot positions Δ***q*** = ***q***_*k*+1_ *−* ***q***^*∗*^ and deviations in the CoM state Δ***s*** = ***s***_*k*_ *−* ***s***^*∗*^ (*k* is the step number). We derived the mapping between Δ***q*** and Δ***s*** at midstance, which was defined as 50% of the step cycle, to be consistent with previous studies and because it was early enough in the gait cycle to allow sufficient time for changes in foot placement by the swing limb [6] [12] [28]. We first estimated this relationship by computing the Jacobian matrix (**J**) during the step cycle that mapped the discrete change in state Δ***s*** to the change in foot position Δ***q*** (Eqn.3 - 4). We assumed left-right symmetry so that the foot positions and the CoM state were mirrored about the sagittal plane [6] [29].

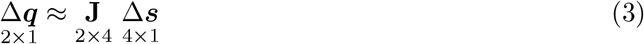

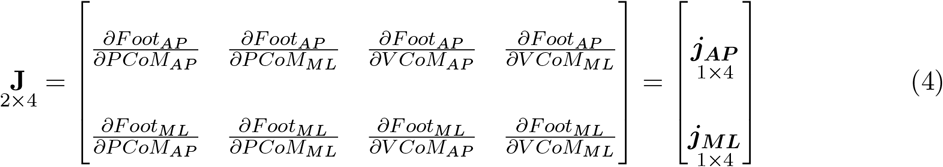

Given that **J** is not a full-rank matrix, and maps from a higher (rank = 4) to a lower (rank = 2) dimension, it has a null space. The null space contains the set of vectors that define the directions along which deviations in CoM state would not affect foot placement. We further defined the first row of **J** matrix to be ***j***_***AP***_ and the second row to be ***j***_***ML***_ as they define how deviations in CoM state influence foot placement in the anteroposterior direction and mediolateral direction, respectively.

### 2.6 Singular Value Decomposition of Jacobian Matrix

The Jacobian matrix can be considered a form of a ”state transition matrix” that reflects the strength and directions of output responses (i.e., changes in foot placements) to inputs (i.e., changes in CoM state) in particular directions in this linearized analysis. Singular value decomposition of the Jacobian, therefore, can estimate the sensitivity of foot placement to changes in CoM state. Importantly, as the Jacobian matrix is not full rank, it maps from higher dimensional changes in CoM state to lower dimensional changes in foot placement. Singular value decomposition can thus determine the changes in CoM state that would produce no changes in foot placement (the null space of the Jacobian). Therefore, we performed singular value decomposition on ***j***_***AP***_ and ***j***_***ML***_ (Eqn. 5) to find their null spaces, determine in which direction the control of foot placement was the most sensitive to deviations in CoM state, and determine the sensitivity of foot placement control along that direction for each individual.

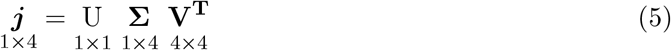

Here, the rank 1, 1 × 4 matrices ***j***_***AP***_ and ***j***_***ML***_ were decomposed as the product of a 1 × 1 matrix U, a 1 × 4 rectangular diagonal gain matrix **Σ**, and a 4 × 4 orthogonal matrix **V**, respectively. The first right singular vector of the Jacobian, ***v***_**1**_, defined the direction along which foot placement was most sensitive to deviations in CoM state. The last three singular vectors (***v***_**2**_, ***v***_**3**_, ***v***_**4**_) defined the null space directions along which deviations in CoM state would not affect the foot placement. The singular values of the gain matrix (Σ) indicated the sensitivity of foot placement to deviations in CoM state along the direction defined by ***v***_**1**_.

### 2.7 Statistical Analysis

Our objective was to determine whether the mapping between CoM state and subsequent foot placement differed between unperturbed and perturbed gait. We combined the data from all participants and used mixed-effects regression to determine the portion of the Jacobian that was consistent across participants (fixed effects) as well as random effects that account for the variability in elements of the Jacobian across participants. We compared the ability of three models to explain anteroposterior and mediolateral foot positions during perturbed walking (Table 1): 1) a linear model derived from unperturbed walking (Model 1, Eqn. 6); 2) a linear model derived from both perturbed steps and unperturbed steps (Model 2, Eqn. 7), and (3) a piecewise linear model derived from both perturbed steps and unperturbed steps (Model 3, Eqn. 8). For Models 2 and 3, we derived foot placement mappings using both the perturbed steps and an equal number of unperturbed steps because a prior study found that foot placement mapping coefficients for unperturbed and backward perturbed walking was similar [6]. Combining step types allowed us to identify a single mapping capable of explaining responses to both internally-generated and external perturbations. We derived a piecewise linear mapping with one breakpoint (Model 3, Eqn. 8) to test for directional differences in responses to increases and reductions in belt speed. We chose this piecewise linear model because there is evidence that people rely on different balance correcting strategies to recover from forward versus backward losses of balance [3] [4] [20] [30].

**Table 1.** Model description for foot placement mappings

| Model Description | Model |  |
| --- | --- | --- |
| Linear mapping derived from unperturbed steps (Model 1) | $\Delta \mathbf{q}_{k+1}^T = \mathbf{J}^1 \Delta \mathbf{s}_k^T$ | (6) |
| Linear mapping derived from both perturbed steps unperturbed steps (Model 2) | $\Delta \mathbf{q}_{k+1}^T = \mathbf{J}^2 \Delta \mathbf{s}_k^T$ | (7) |
| A piecewise linear regression model derived from both perturbed steps and unperturbed steps (Model 3) | $\Delta \mathbf{q}_{k+1}^T = \begin{cases} \mathbf{J}^3 \Delta \mathbf{s}_k^T & \text{if } \Delta VCoM_{AP} > 0 \\ \mathbf{J}^4 \Delta \mathbf{s}_k^T & \text{if } \Delta VCoM_{AP} < 0 \end{cases}$ | (8) |

We used the AIC to determine the most parsimonious model to explain variance in foot placement (Eqn. 9) [31].

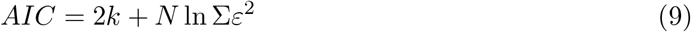

Here, *k* is the number of estimated parameters, *N* is the number of data points, *ε* is the prediction error between the predicted and actual data. We selected the model with the lowest AIC as the best model.

We also determined if the foot placement mapping differed between perturbed and unperturbed walking by comparing the regression coefficients of the foot placement mapping derived from perturbed walking and those derived from unperturbed walking. Lastly, we determined whether the values of the gain matrix from singular value decomposition that indicated the sensitivity of foot placement control in response to deviations in CoM state differed between unperturbed walking and perturbed walking. We used paired sample t-test if the variables were normally distributed; otherwise, we used Wilcoxon rank-sum test. We used the Shapiro-Wilk Test to test the normality. Significance was set at p*<*0.05.

## 3 Results

### 3.1 Foot placement mapping during unperturbed walking

Both anteroposterior and mediolateral foot position relative to the trailing limb varied from step to step during unperturbed walking (Figure 2A, grey points). Foot position in the anteroposterior direction was explained by a model which included CoM displacement and velocity in both anteroposterior and mediolateral direction with the following form (mean *±* standard error) which had an R^2^ of 0.38:

**Figure 2.**
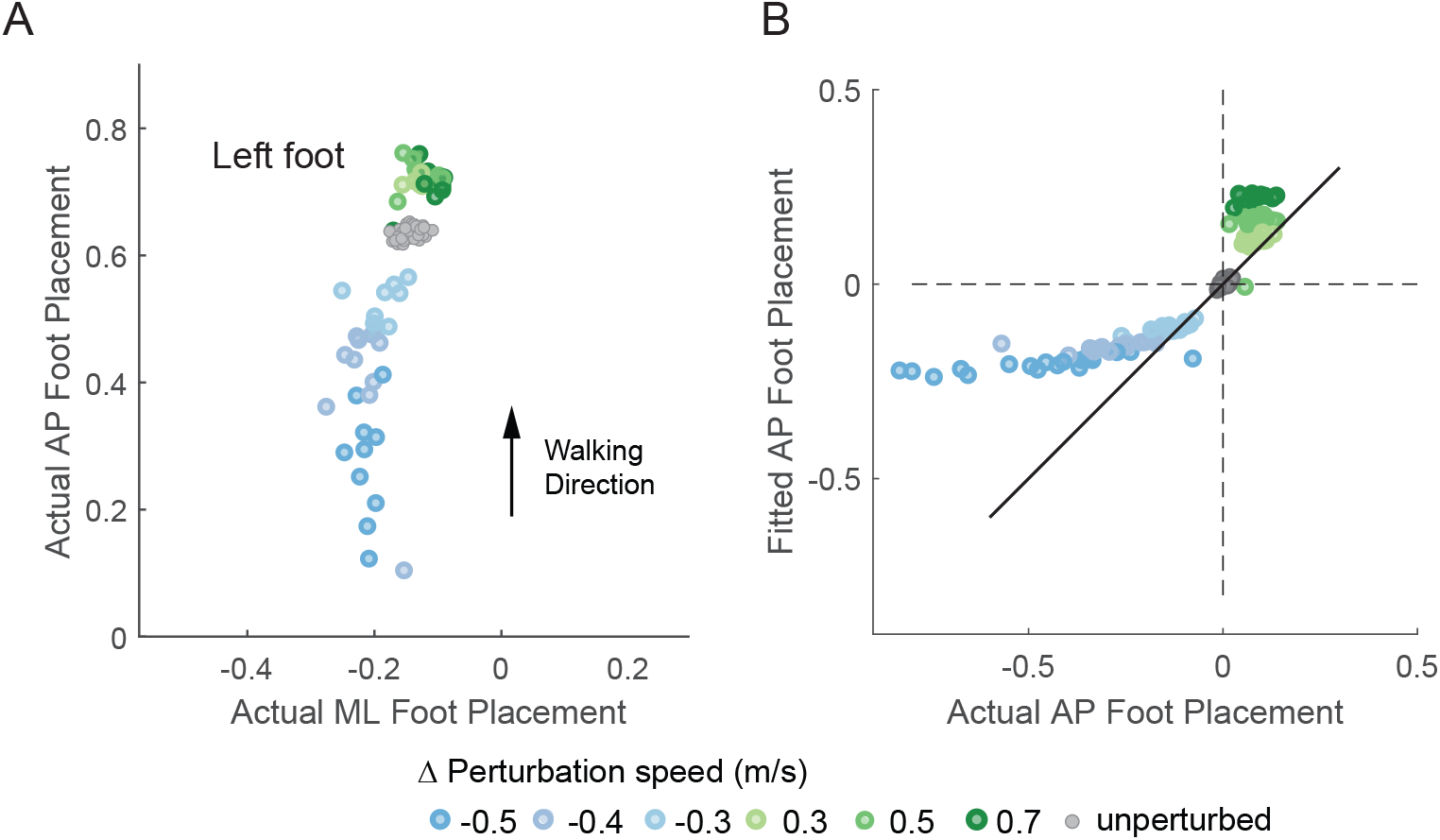
Scatter plots showing the left foot placement during unperturbed walking and following perturbations for a representative participant. Colored dots indicate foot placement following increasing perturbations (blue to green). Gray dots represent foot placement during unperturbed walking. (A) Left foot placement relative to the right perturbed trailing stance foot during unperturbed steps and perturbed steps. (B) Actual foot placement v. fitted foot placement in the anteroposterior direction during both unperturbed and perturbed walking using the mapping derived from unperturbed steps.

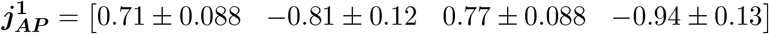

Thus, a larger forward displacement of the CoM and larger forward velocity at midstance were associated with a longer step while a larger lateral CoM displacement and larger lateral velocity at midstance were associated with a shorter step. Foot position in the mediolateral direction was positively associated with CoM displacement and velocity in the mediolateral direction at midstance and negatively associated with CoM velocity in the anteroposterior direction which had an R^2^ of 0.74:

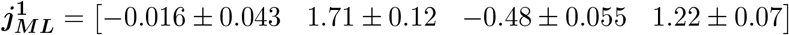

### 3.2 Foot placement mapping during perturbed walking

The mapping between foot position and CoM state at midstance during unperturbed waking did not generalize to foot positions following perturbations based on visual inspection of the predictions from the unperturbed model (Figure 2). In both mediolateral and anteroposterior directions, we found that a piecewise linear model best explained the variance in foot placement as evidenced by the lower AIC values (Table 2). Following forward perturbations, a larger forward displacement and larger forward velocity of the CoM at midstance were associated with a longer step while a larger lateral CoM velocity and larger lateral velocity at midstance were associated with a shorter step.

**Table 2.**
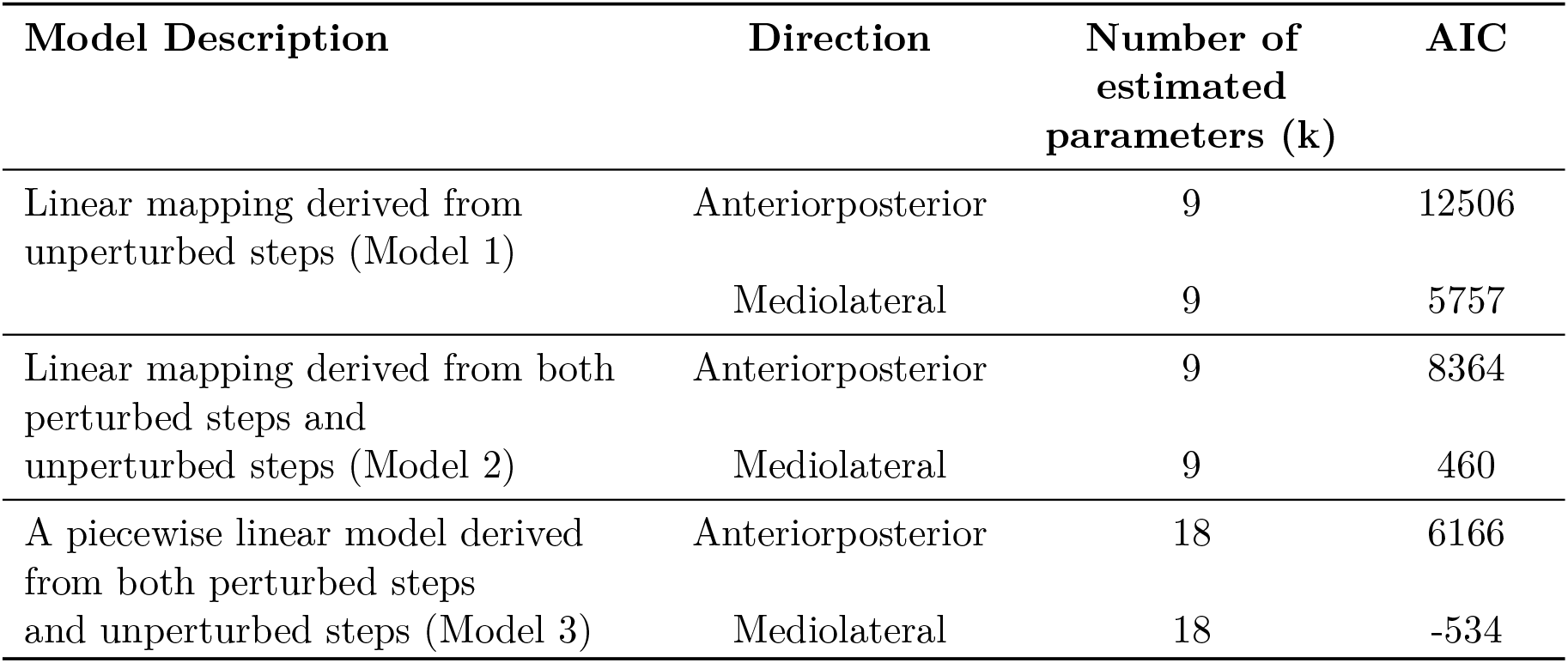
Model selection metrics based on AIC. Lower AIC values are indicative of better models.

| Model Description | Direction | Number of<br>estimated<br>parameters (k) | AIC |
| --- | --- | --- | --- |
| Linear mapping derived from unperturbed steps (Model 1) | Anteriorposterior | 9 | 12506 |
|  | Mediolateral | 9 | 5757 |
| Linear mapping derived from both perturbed steps and unperturbed steps (Model 2) | Anteriorposterior | 9 | 8364 |
|  | Mediolateral | 9 | 460 |
| A piecewise linear model derived from both perturbed steps and unperturbed steps (Model 3) | Anteriorposterior | 18 | 6166 |
|  | Mediolateral | 18 | -534 |

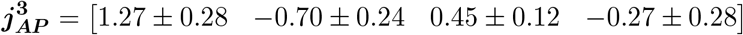

On the other hand, following backward perturbations, a larger backward displacement and larger backward velocity of CoM were associated with a shorter step while a larger lateral CoM displacement and larger medial velocity at midstance were associated with a shorter step.

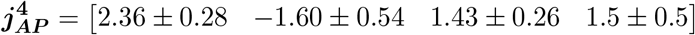

In the mediolateral direction, a larger lateral CoM velocity and displacement at midstance were associated with a wider step for both forward and backward perturbations. A larger forward CoM displacement and velocity were associated with a narrower step following forward perturbations. For backward perturbations, a larger backward CoM displacement and smaller backward CoM velocity were associated with a narrower step.

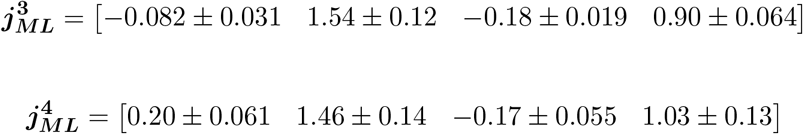

Several features of the anteroposterior foot placement mappings differed depending on the dataset for which they were derived (Figure 3A). Coefficient estimates for each individual were computed by summing the random effects and the fixed effects from each mixed effect model. The coefficients for Δ*PCoM*_*AP*_ derived from backward perturbations were greater than those derived from forward perturbations (t(12) = 4.3, p = 0.0011) and unperturbed walking (t(12) = 5.6, p = 0.0001). Similarly, the coefficients for Δ*V CoM*_*AP*_ derived from backward perturbations were greater than those derived from forward perturbations (t(12) = 2.4, p = 0.034) and unperturbed walking (t(12) = 3.6, p = 0.0037). This suggests that, for a fixed magnitude deviation in CoM state, changes in foot placement were larger in response to backward versus forward perturbations. The coefficients for Δ*V CoM*_*ML*_ were greater when derived from forward perturbations than unperturbed walking (t(12) = 3.5, p = 0.0043). The coefficients for Δ*V CoM*_*ML*_ derived from backward perturbations were also greater than those derived from forward perturbations (t(12) = 3.1, p = 0.0093) and unperturbed walking (t(12) = 5.0, p = 0.0003) and were generally positive while those derived from forward perturbations and unperturbed walking were generally negative. This suggests that a fixed magnitude of deviation in lateral CoM velocity would result in a longer step during backward perturbations but a shorter step during unperturbed walking and forward perturbations.

**Figure 3.**
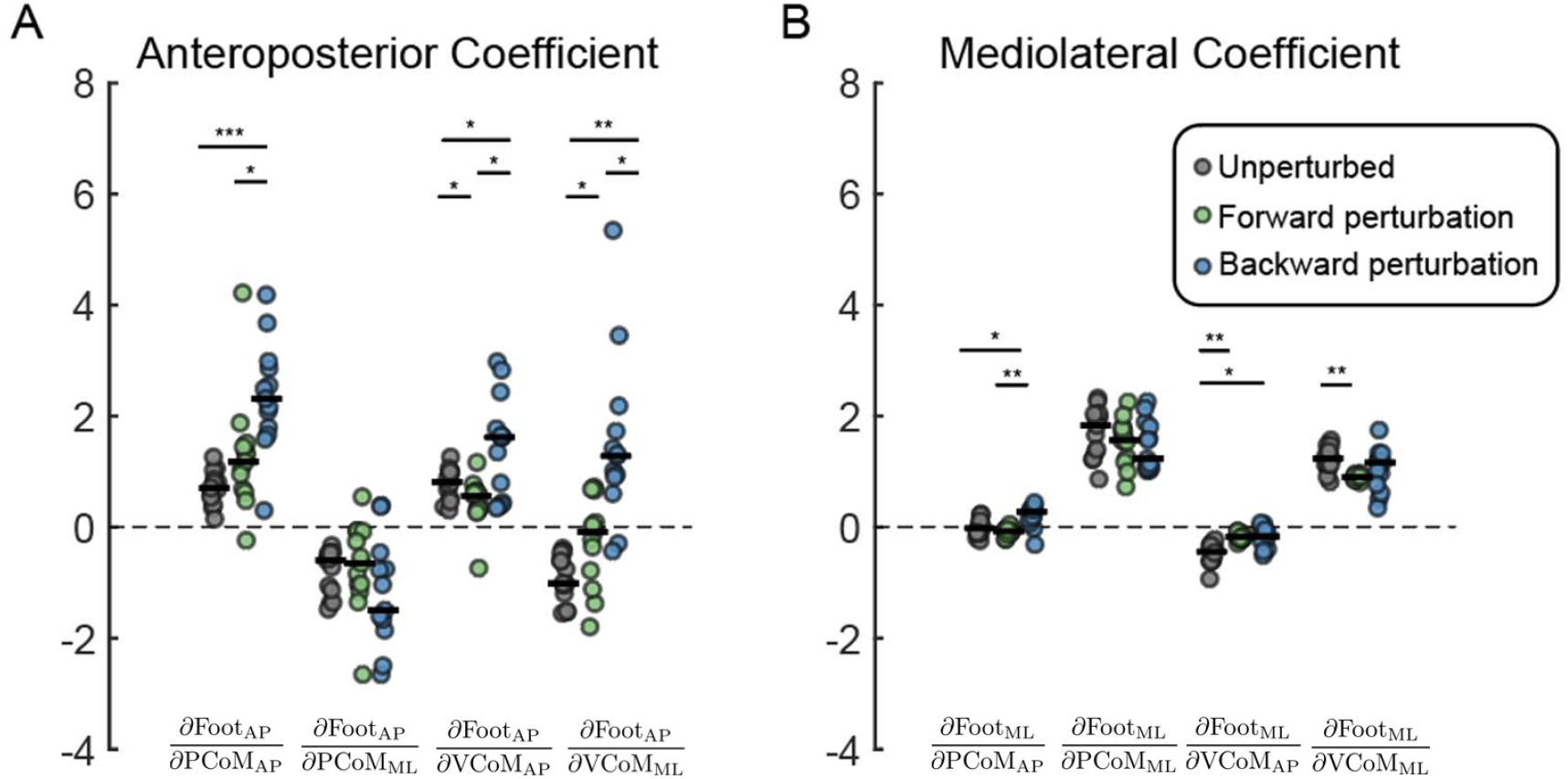
The estimated coefficients of the foot placement model in the anteroposterior direction (A) and mediolateral direction (B) with respect to CoM state at midstance. Coefficient estimates were computed by summing the random effects and the fixed effects from each mixed effect model. Black horizontal lines indicate the median coefficient estimates across participants. Gray: estimates from unperturbed walking (Model 1), Green: estimates from piecewise linear model for forward perturbations (Model 3), Blue: estimates from piecewise linear model for backward perturbations (Model 3). Dots represented individual estimates of coefficients (*p*<*0.05, **p*<*0.001,***p*<*0.0001).

The mediolateral foot placement mapping derived from perturbed walking differed from that derived from unperturbed walking (Figure 3B). The coefficients for Δ*PCoM*_*AP*_ derived from backward perturbations were higher than those from unperturbed walking (t(12) = 3.82, p = 0.0024) and forward perturbations (t(12) = 4.79, p = 0.0004). The coefficients for Δ*V CoM*_*AP*_ derived from unperturbed walking were more negative than derived from forward perturbations (t(12) = -5.3, p = 0.0002) and backward perturbations (t(12) = -4.1, p = 0.0014). Lastly, the coefficients for Δ*V CoM*_*ML*_ derived from forward perturbations were less than those derived from unperturbed walking (t(12) = -5.3, p = 0.0002).

Although participants experienced many perturbations over the course of the experiment, we did not observe learning effects as measured by their responses to the perturbations. To assess the potential for learning, we compared the distance from the CoM to the rear edge of the base of support and also compared the CoM velocity in the anteroposterior direction at the time of foot strike after the first and last perturbations for each level of treadmill speed change [32]. There were no differences in these measures between the first and last perturbations (CoM position: p = 0.25; CoM velocity: p = 0.20) indicating that participants responded similarly to the perturbations throughout the experiment.

### 3.3 Singular Value Decomposition of Foot Placement Mappings

#### 3.3.1 Task space vectors for anteroposterior foot placement mapping matrix

Singular value decomposition provided a direct assessment of the null space of **J**, and the directions along which future foot placement Δ***q*** was the most sensitive to changes in CoM state Δ***s*** (Figure 4A-F blue arrows). We first performed singular value decomposition on the Jacobian matrix obtained for unperturbed walking 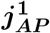, forward perturbations 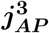, and backward perturbations 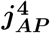 in the anteroposterior direction. During unperturbed walking 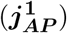, the largest foot placement changes were associated with deviations in CoM displacement and velocity that were directed anteriorly and medially (Figure 4A-D). This was consistent with our interpretation in Section 3.1 that a larger forward displacement of the CoM and larger forward velocity at midstance were associated with a longer step, while a larger lateral CoM displacement and larger lateral velocity at midstance were associated with a shorter step. Following forward perturbations 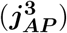, people generally made the largest adjustment in foot placement in response to deviations in CoM displacement and velocity that were directed anteriorly and medially (Figure 4E-H). However, it is important to note that there was large inter-subject variability in response to deviations in CoM velocity in this case (Figure 4G). Unlike the unperturbed and forward perturbation conditions, during the backward perturbations 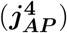 the largest changes in foot placement were associated with posterior/lateral deviations of CoM displacement coupled with posterior/medial deviations in CoM velocity (Figure 4I-L). The direction for deviations in CoM velocity was different from unperturbed steps and forward perturbations. Thus, these results suggest that changes in foot placement were direction-dependent in response to forward and backward perturbations in terms of CoM velocity, but the mapping remained relatively invariant in terms of CoM displacement.

**Figure 4.**
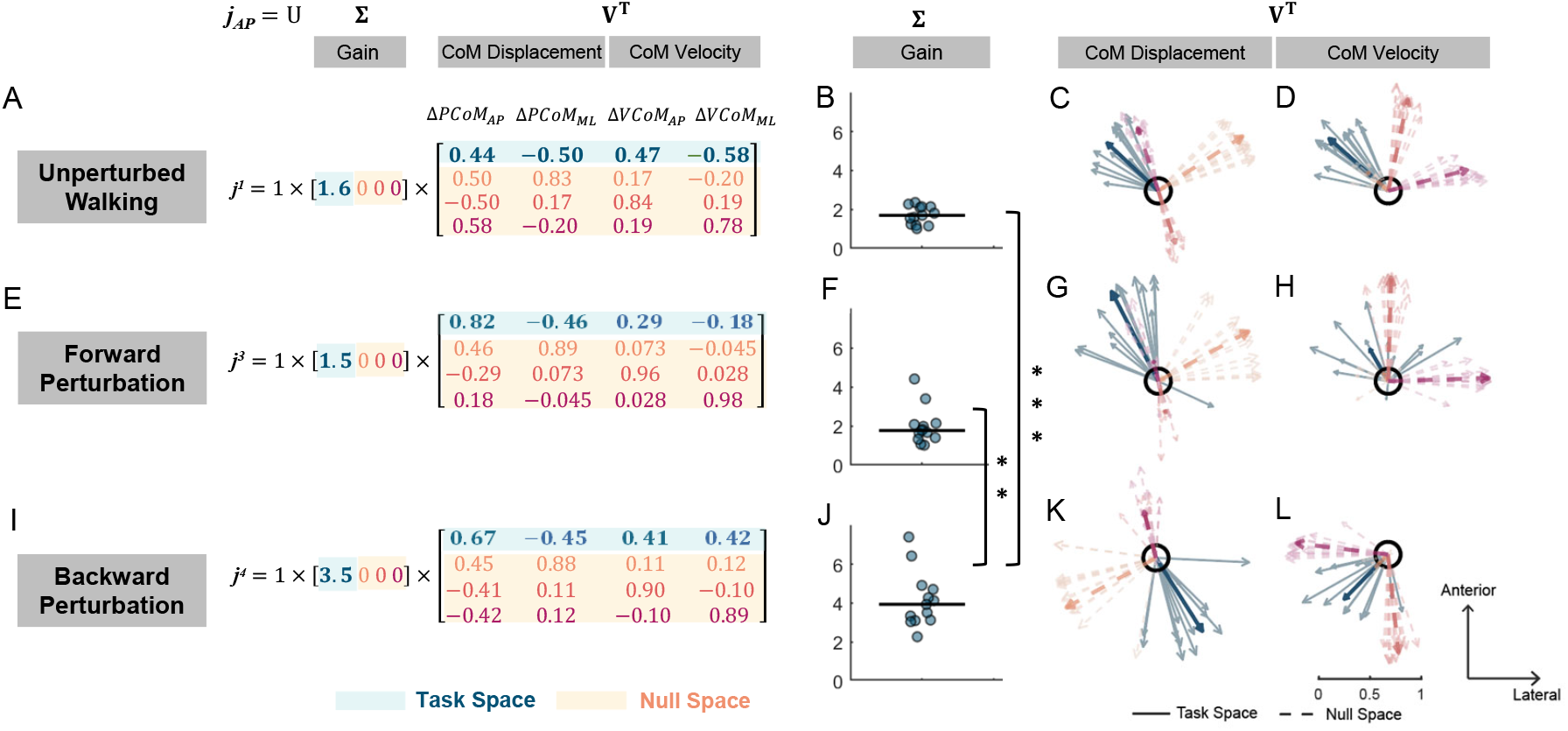
Visualization of singular value decomposition of the anteroposterior foot placement mapping matrix derived from unperturbed steps, forward perturbation, and backward perturbation steps. Left panel shows singular value decomposition on the mean foot placement mapping matrix derived from unperturbed steps (A), forward perturbation (E), and backward perturbation steps (I). Gain obtained from singular value decomposition on the foot placement mapping for unperturbed steps (B), forward perturbation (F), and backward perturbations (J) for each individual (dot) and median across participants (black line). (**p*<*0.001,***p*<*0.0001). Right singular vectors related with ΔCoM displacement derived during steady-state walking (C), during forward perturbation (G), during backward perturbation (K). Right singular vectors related with ΔCoM velocity derived from mapping coefficients during steady-state walking (D), during forward loss of balance (H), during backward loss of balance (L). Light colored arrows indicate right singular vectors for each individual. Note that solid arrows indicate the first right singular vector (task space vectors) while dash lines indicate the last three singular vectors (null space vectors). Dark colored arrows indicate right singular vectors computed from the mean foot placement mapping matrix.

#### 3.3.2 Null space vectors for anteroposterior foot placement mapping matrix

Deviations in CoM state along the last three singular vectors (null space vectors) would not affect the foot placement. The orientations of null space vectors were similar for unperturbed walking and forward and backward perturbations. During both unperturbed and perturbed steps, deviations in CoM displacement that were directed anteriorly and laterally would not affect foot placement position (Figure 4 orange arrows). Deviations in CoM velocity in the lateral direction would also not affect foot placement position (Figure 4 red arrows). Lastly, deviations in CoM velocity directed anteriorly coupled with deviations in CoM displacement directed posteriorly would not affect foot placement position (Figure 4 pink arrows).

#### 3.3.3 Gain values for anteroposterior foot placement mapping matrix

Singular value decomposition of the anteroposterior foot placement mapping revealed higher control gain during backward perturbation than unperturbed walking and forward perturbation. The gain obtained for backward perturbations was higher than the gain obtained for unperturbed (Z = 4.2, p *<* 0.0001) and forward perturbation (p = 0.0003; Figure 4B, F, J). These results indicated that foot placement was more sensitive to the changes in CoM state and may be more tightly controlled during backward perturbation than forward perturbation or unperturbed walking.

#### 3.3.4 Task space vectors for mediolateral foot placement mapping matrix

Similarly, we performed singular value decomposition on the Jacobian matrix obtained for unperturbed walking 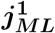, forward perturbations 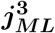, and backward perturbations 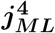 in the mediolateral direction (Figure 5). During both unperturbed walking and perturbed walking, a larger lateral displacement and velocity at midstance were associated with a wider step (Figure 5 blue arrows). This was consistent with our results in Section 3.1 that a larger lateral displacement of the CoM and larger lateral velocity at midstance were associated with a longer step.

**Figure 5.**
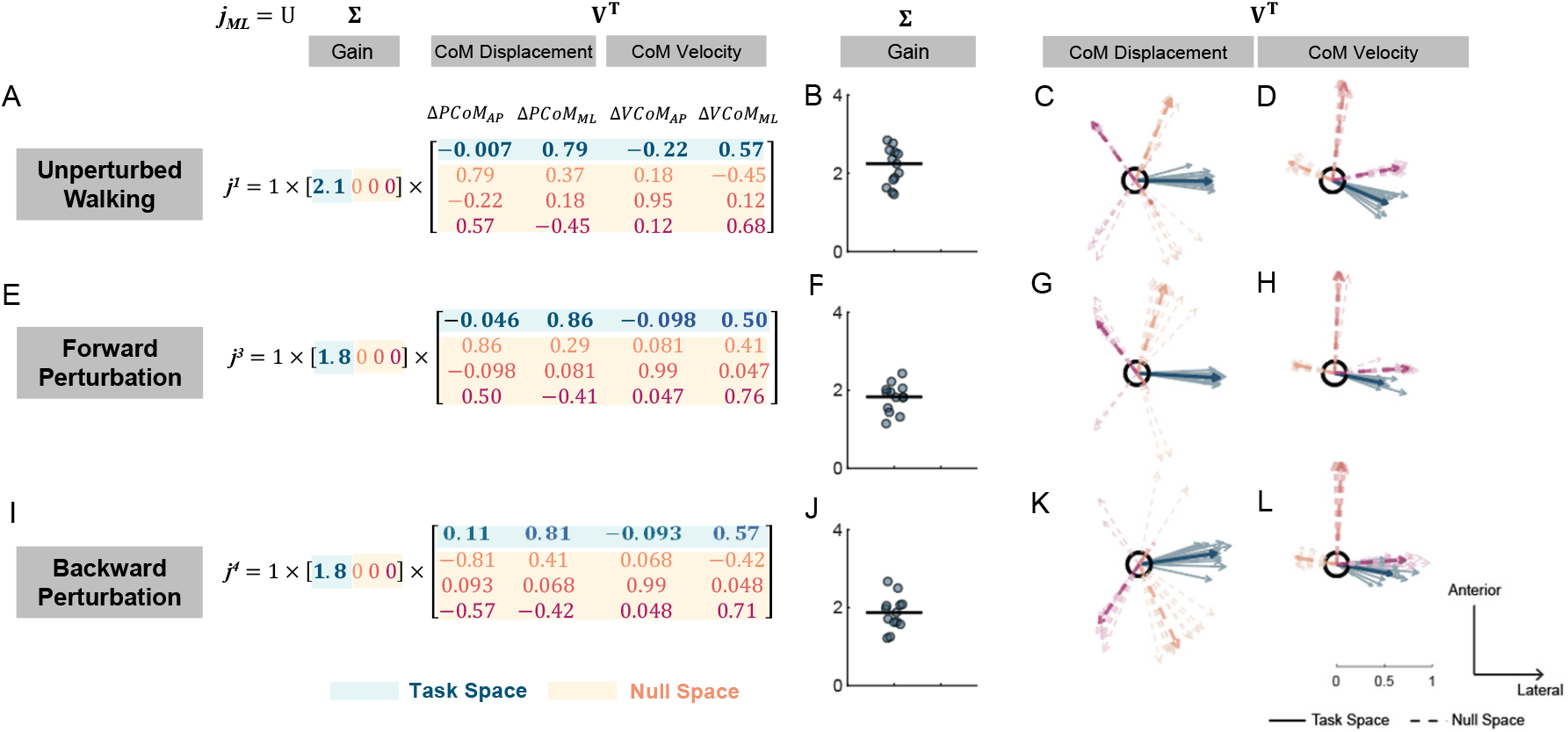
Visualization of singular value decomposition of the mediolateral foot placement mapping matrix derived from unperturbed steps, forward perturbation, and backward perturbation steps. Left panel shows singular value decomposition on the mean foot placement mapping matrix derived from unperturbed steps (A), forward perturbation (E), and backward perturbation steps (I). Gain obtained from singular value decomposition on the foot placement mapping for unperturbed steps (B), forward perturbation (F), and backward perturbations (J) for each individual (dot) and median across participants (black line). Right singular vectors related with ΔCoM displacement derived during steady-state walking (C), during forward perturbation (G), during backward perturbation (K). Right singular vectors related with ΔCoM velocity derived from mapping coefficients during steady-state walking (D), during forward loss of balance (H), during backward loss of balance (L). Light colored arrows indicate right singular vectors for each individual. Note that solid arrows indicate the first right singular vector (task space vectors) while dash lines indicate the last three singular vectors (null space vectors). Dark colored arrows indicate right singular vectors computed from the mean foot placement mapping matrix.

#### 3.3.5 Null space vectors for mediolateral foot placement mapping matrix

The directions of null space vectors were similar for unperturbed walking and forward but not for backward perturbations. For unperturbed walking and forward perturbations, deviations in CoM displacement that were directed anteriorly and laterally did not affect foot placement position (Figure 5C, D, G, H orange arrows). Deviations in CoM velocity in the fore-aft direction also did not affect mediolateral foot placement position (Figure 5C, D, G, H red arrows). Deviations in CoM velocity directed laterally coupled with deviations in CoM displacement directed anteriorly and medially did not affect foot placement position (Figure 5C, D, G, H pink arrows). Following backward perturbations, deviations in CoM displacement that were directed posteriorly and medially did not affect foot placement position (Figure 5K, L orange arrows). Deviations in CoM velocity in the fore-aft direction also did not affect mediolateral foot placement position (Figure 5K, L red arrows).

Deviations in CoM velocity directed laterally coupled with deviations in CoM displacement directed posteriorly and medially would not affect foot placement position (Figure 5K, L pink arrows).

#### 3.3.6 Gain values for mediolateral foot placement mapping matrix

Lastly, singular value decomposition on mediolateral foot placement mapping found similar gain during unperturbed walking, following forward and backward perturbations (p *>* 0.05; Figure 5B, F, J). Such results indicated that sensitivity of mediolateral foot placement to the changes in CoM state was similar during unperturbed walking and forward or backward perturbations.

## 4 Discussion

Our study’s primary objective was to determine if the mapping between changes in CoM state and changes in foot placement found during steady-state, unperturbed walking explained changes in foot placement in response to imposed perturbations. We found that the mapping derived from the natural variability of foot placement during steady-state walking could not explain patterns of foot placement in response to perturbations

(Figure 2B). Instead, a mapping that accounted for differences in responses to forward versus backward perturbations best explained foot placement variance during perturbed steps (Table 2). In addition, we found that foot placement was more sensitive to the changes in CoM state and more tightly correlated with backward perturbations than forward perturbation. Overall, our results demonstrate that a mapping that accounted for directional differences emerges when people adjust their foot placement in response to forward and backward perturbations.

The foot placement mapping during unperturbed walking in neurotypical participants was similar to that previously reported for young adults despite the fact that our population was, on average, older [12]. Our derived foot placement mappings explained *∼*60% of the variance in foot placement in the mediolateral direction and *∼*40% of the variance in the anteroposterior direction at midstance, which is comparable with prior work [12] [15]. In the fore-aft direction, more lateral deviation of CoM displacement and CoM velocity at midstance was associated with a shorter step while a more forward deviation of CoM displacement and CoM velocity was associated with a longer step. In the mediolateral direction, more lateral deviation of CoM displacement and velocity was associated with a more lateral step. In both directions, people stepped in the direction of the CoM deviation. Such association between deviation in CoM state and foot placement could be attributed, in part, to passive dynamics of the swing leg and active control of foot placement to maintain balance [9] [16]. Additionally, as in the neurotypical young population, the coefficient of determination at midstance was higher for mediolateral deviations in foot placement than fore-aft deviations, indicating that people may adopt a tighter control their foot placement in the mediolateral direction than in the fore-aft direction.

We hypothesized that a mapping that accounted for the differences in response to forward versus backward disturbances would better explain the variance in foot placement than a linear mapping derived from unperturbed walking. Consistent with this hypothesis, we found that the foot placement mapping differed between forward versus backward perturbations. For instance, changes in foot placement in the anteroposterior direction were more sensitive to changes in fore-aft CoM displacement and velocity at midstance following backward perturbations than forward perturbations. The discrepancy in foot placement mapping between forward and backward perturbations may result from the fact that people rely more on modulation of ankle torque in the perturbed limb during forward perturbations than they do during backward perturbations [4] [5]. Shifting the center of pressure forward by activating the ankle plantar flexors during the stance phase in which forward perturbations occur could help people to generate backward moment about body CoM to reduce the forward rotation of the body. As a result, a smaller backward moment needs to be generated about the body’s CoM at the next foot placement and less foot placement deviation from the nominal trajectory was needed in response to forward perturbations than backward perturbations.

The mediolateral foot placement mapping derived from unperturbed walking also differed from the mapping derived from perturbed walking. Similar to what was observed with foot placement in the anteroposterior direction, these results indicate that the mapping between CoM state and foot placement observed during unperturbed walking does not generalize to perturbed walking. These results may indicate that our nervous system adjusts the control strategies following perturbations to generate appropriate corrective responses to maintain balance. This difference in control between steady-state and perturbed walking may reflect a shift from more spinally-mediated control to control by brainstem or cortical circuits responsible for balance control [33] [34] [35]. For example, treadmill accelerations and decelerations which were similar to the perturbation paradigm used in this current study induced long-latency stretch reflexes in calf muscles that are thought to be mediated by supraspinal structures [36]. Therefore, analysis of unperturbed walking is insufficient to infer control strategies responsible for recovering from losses of balance.

The use of singular value decomposition extended our interpretations of foot placement control strategies beyond what could be inferred solely from the derived foot placement mappings. Performing singular value decomposition on the Jacobian matrix has been widely used for analyzing and designing control systems [37]. In our analysis, we applied the decomposition to the experimental Jacobian matrices to obtain the direction along which changes in foot placement was most sensitive to changes in CoM state and the sensitivity (gain) along that direction. We found that both the direction and gain were similar for unperturbed steps and following forward perturbations. In contrast, the direction and gain were different following backward perturbations. This suggests that foot placement control strategies following backward perturbations were different from strategies during unperturbed and following forward perturbations. Particularly, the gain for backward perturbations was greater than unperturbed and forward perturbations, indicating higher sensitivity to deviations in CoM state following backward perturbations and, we speculate, tighter control of foot placement to correct for such deviations in CoM state compared to unperturbed and following forward perturbations.

Other stabilization strategies aside from foot placement, such as modulating the ankle push-off, also play an important role in maintaining balance [5] [15] [30] [38] [39]. We previously demonstrated that neurotypical participants coordinate both their leading and trailing limb to restore balance in response to forward loss of balance [5]. Kim and Collins [28] derived a controller that used both foot placement and ankle push-off impulse to stabilize a biped in the sagittal plane when negotiating through random changes of the ground’s height during walking. Therefore, future studies may investigate how different balance recovery strategies coordinate together following the deviation in body’s state and whether such coordination may explain the difference in foot placement mapping following the forward and backward perturbations.

Although we used CoM state as the predictor to derive the foot placement mapping, it is uncertain if CoM state provides the best predictive value. Other studies have used the swing leg state at the swing initiation [14], the stance leg state [13], or the ankle state [40] to construct predictive models that describe how humans control balance during walking or running. Future studies should perform a more comprehensive model comparison to determine the best set of predictors to explain foot placement control.

It also remains unclear to what extent passive dynamics versus active control contribute to the observed associations between CoM state and foot placement. For example, an open-loop stable 2D model showed that 80% of the variance in foot position could be explained by CoM state in the fore-aft direction at midstance [16]. One primary objective of our study was to derive the foot placement mapping during relatively large perturbations that required reactive responses to avoid falls. To our knowledge, no studies have examined the role of passive dynamics during balance corrections for perturbed walking. Given the inability of mappings derived from unperturbed walking to explain the variance in foot placement in the current study, this may suggest a larger contribution from active control in response to external perturbations. In addition, the previously examined 2D bipedal model did not consider the inertial properties of the swing limb or consider control of the torso that helps to maintain an upright posture [16]. Thus, a more complex model with segment inertias [41] may be necessary to untangle the relative contribution of passive dynamics and active control to the correlation between body’s state and foot placement and draw inference about how people use sensory feedback information to generate corrective response.

## 5 Data Availability

All data can be retrieved from:

https://osf.io/gv5tq/?view_only=858243326d374cd3ba6ddd157195d02f

## 6 Acknowledgments

We would like to thank Aram Kim and Isaiah Lachica for their help with reviewing the figures and the manuscript. This work was supported by the Eunice Kennedy Shriver National Institute of Child Health and Human Development of the National Institutes of Health under Award Number R01HD091184.

## References

1. Bruijn SM, Meijer OG, Beek P, van Dieen JH. Assessing the Stability of Human Locomotion: A Review of Current Measures. Journal of The Royal Society Interface. 2013;10(83):20120999–20120999. doi:10.1098/rsif.2012.0999.

2. Redfern MS, Schumann T. A Model of Foot Placement during Gait. Journal of Biomechanics. 1994;27(11):1339–1346. doi:10.1016/0021-9290(94)90043-4.

3. Reimann H, Fettrow T, Thompson ED, Jeka JJ. Neural Control of Balance during Walking. Frontiers in Physiology. 2018;9:1–13. doi:10.3389/fphys.2018.01271.

4. Vlutters M, Van Asseldonk EHF, Van Der Kooij H. Center of Mass Velocity-Based Predictions in Balance Recovery Following Pelvis Perturbations during Human Walking. Journal of Experimental Biology. 2016;219(10):1514–1523. doi:10.1242/jeb.129338.

5. Liu C, McNitt-Gray JL, Finley JM. Impairments in the Mechanical Effectiveness of Reactive Balance Control Strategies during Walking in People Post-Stroke. Frontiers in Neurology. 2022;13.

6. Joshi V, Srinivasan M. A Controller for Walking Derived from How Humans Recover from Perturbations. Journal of the Royal Society Interface. 2019-08-30;16(157):20190027. doi:10.1098/rsif.2019.0027.

7. Mahaki M, Bruijn SM, van Dieën JH. The Effect of External Lateral Stabilization on the Use of Foot Placement to Control Mediolateral Stability in Walking and Running. PeerJ. 2019;7:e7939. doi:10.7717/peerj.7939.

8. Perry JA, Srinivasan M. Walking with Wider Steps Changes Foot Placement Control, Increases Kinematic Variability and Does Not Improve Linear Stability. Royal Society Open Science. 2017;4(9). doi:10.1098/rsos.160627.

9. Rankin BL, Buffo SK, Dean JC. A Neuromechanical Strategy for Mediolateral Foot Placement in Walking Humans. Journal of Neurophysiology. 2014;112(2):374–383. doi:10.1152/jn.00138.2014.

10. Roden-Reynolds DC, Walker MH, Wasserman CR, Dean JC. Hip Proprioceptive Feedback Influences the Control of Mediolateral Stability during Human Walking. Journal of Neurophysiology. 2015;114(4):2220–2229. doi:10.1152/jn.00551.2015.

11. Stimpson KH, Heitkamp LN, Horne JS, Dean JC. Effects of Walking Speed on the Step-by-Step Control of Step Width. Journal of Biomechanics. 2018;68:78–83. doi:10.1016/j.jbiomech.2017.12.026.

12. Wang Y, Srinivasan M. Stepping in the Direction of the Fall: The next Foot Placement Can Be Predicted from Current Upper Body State in Steady-State Walking. Biology Letters. 2014;10(9). doi:10.1098/rsbl.2014.0405.

13. Arvin M, Hoozemans MJM, Pijnappels M, Duysens J, Verschueren SM, van Dieën JH. Where to Step? Contributions of Stance Leg Muscle Spindle Afference to Planning of Mediolateral Foot Placement for Balance Control in Young and Old Adults. Frontiers in Physiology. 2018;9:1–10. doi:10.3389/fphys.2018.01134.

14. Dean JC, Kautz SA. Foot Placement Control and Gait Instability among People with Stroke. Journal of rehabilitation research and development. 2015;52(5):577–590. doi:10.1682/JRRD.2014.09.0207.

15. Jin J, van Dieën JH, Kistemaker D, Daffertshofer A, Bruijn SM. Does Ankle Push-off Correct for Errors in Anterior–Posterior Foot Placement Relative to Center-of-Mass States?; 2023.

16. Patil NS, Dingwell JB, Cusumano JP. Correlations of Pelvis State to Foot Placement Do Not Imply Within-Step Active Control. Journal of Biomechanics. 2019;97. doi:10.1016/j.jbiomech.2019.109375.

17. Dal U, Erdogan T, Resitoglu B, Beydagi H. Determination of Preferred Walking Speed on Treadmill May Lead to High Oxygen Cost on Treadmill Walking. Gait and Posture. 2010-03-01;31(3):366–369. doi:10.1016/j.gaitpost.2010.01.006.

18. Farrens A, Lilley M, Sergi F. Training Propulsion via Acceleration of the Trailing Limb. IEEE Transactions on Neural Systems and Rehabilitation Engineering. 2020-12;28(12):2816–2825. doi:10.1109/tnsre.2020.3032094.

19. García-Pérez MA. Forced-Choice Asymptotic and Small-Sample Properties. Vision Research. 1998-06-01;38(12):1861–1881. doi:10.1016/S0042-6989(97)00340-4.

20. Liu C, Park S, Finley J. The Choice of Reference Point for Computing Sagittal Plane Angular Momentum Affects Inferences about Dynamic Balance. PeerJ. 2022;10:e13371. doi:10.7717/peerj.13371.

21. Liu C, Macedo LD, Finley JM. Conservation of Reactive Stabilization Strategies in the Presence of Step Length Asymmetries During Walking. Frontiers in Human Neuroscience. 2018;12.

22. Havens KL, Mukherjee T, Finley JM. Analysis of Biases in Dynamic Margins of Stability Introduced by the Use of Simplified Center of Mass Estimates during Walking and Turning. Gait & Posture. 2018;59:162–167. doi:10.1016/j.gaitpost.2017.10.002.

23. Song J, Sigward S, Fisher B, Salem GJ. Altered Dynamic Postural Control during Step Turning in Persons with Early-Stage Parkinson’s Disease. Parkinson’s Disease. 2012;2012:386962. doi:10.1155/2012/386962.

24. Kurz MJ, Arpin DJ, Corr B. Differences in the Dynamic Gait Stability of Children with Cerebral Palsy and Typically Developing Children. Gait and Posture. 2012;36(3):600–604. doi:10.1016/j.gaitpost.2012.05.029.

25. Reisman DS, Wityk R, Silver K, Bastian AJ, Manuscript A. Split-Belt Treadmill Adaptation Transfers to Overground Walking in Persons Poststroke. Physical Therapy. 2009;23(7):735–744. doi:10.1177/1545968309332880.Split-Belt.

26. Winter DA. Biomechanics and Motor Control of Human Movement. vol. 2nd; 2009.

27. Buurke TJW, Liu C, Park S, den Otter R, Finley JM. Maintaining Sagittal Plane Balance Compromises Frontal Plane Balance during Reactive Stepping in People Post-Stroke. Clinical Biomechanics. 2020-12-01;80:105135. doi:10.1016/j.clinbiomech.2020.105135.

28. Kim M, Collins SH. Once-Per-Step Control of Ankle Push-Off Work Improves Balance in a Three-Dimensional Simulation of Bipedal Walking. IEEE Transactions on Robotics. 2017-04-01;33(2):406–418. doi:10.1109/TRO.2016.2636297.

29. Ankaralı MM, Sefati S, Madhav MS, Long A, Bastian AJ, Cowan NJ. Walking Dynamics Are Symmetric (Enough). Journal of The Royal Society Interface. 2015;12(108):20150209. doi:10.1098/rsif.2015.0209.

30. Pijnappels M, Bobbert MF, van Dieën JH. Push-off Reactions in Recovery after Tripping Discriminate Young Subjects, Older Non-Fallers and Older Fallers. Gait & Posture. 2005-06;21(4):388–394. doi:10.1016/j.gaitpost.2004.04.009.

31. Akaike H. Likelihood of a Model and Information Criteria. Journal of Econometrics. 1981-05-01;16(1):3–14. doi:10.1016/0304-4076(81)90071-3.

32. Wang TY, Bhatt T, Yang F, Pai YCC. Adaptive Control Reduces Trip-Induced Forward Gait Instability among Young Adults. Journal of Biomechanics. 2012;45(7):1169–1175. doi:10.1016/j.jbiomech.2012.02.001.

33. Deliagina TG, Orlovsky GN, Zelenin PV, Beloozerova IN. Neural Bases of Postural Control. Physiology. 2006-06;21(3):216–225. doi:10.1152/physiol.00001.2006.

34. Finley JM, Dhaher YY, Perreault EJ. Acceleration Dependence and Task-Specific Modulation of Short- and Medium-Latency Reflexes in the Ankle Extensors. Physiological Reports. 2013;1(3). doi:10.1002/phy2.51.

35. Solopova IA, Kazennikov OV, Deniskina NB, Levik YS, Ivanenko YP. Postural Instability Enhances Motor Responses to Transcranial Magnetic Stimulation in Humans. Neuroscience Letters. 2003-01-30;337(1):25–28. doi:10.1016/S0304-3940(02)01297-1.

36. Sloot LH, Van Den Noort JC, Van Der Krogt MM, Bruijn SM, Harlaar J. Can Treadmill Perturbations Evoke Stretch Reflexes in the Calf Muscles? PLoS ONE. 2015-12-01;10(12). doi:10.1371/journal.pone.0144815.

37. Klema V, Laub A. The Singular Value Decomposition: Its Computation and Some Applications. IEEE Transactions on Automatic Control. 1980-04;25(2):164–176. doi:10.1109/TAC.1980.1102314.

38. Afschrift M, Groote FD, Jonkers I. Similar Sensorimotor Transformations Control Balance during Standing and Walking. PLOS Computational Biology. 2021;17(6):e1008369. doi:10.1371/journal.pcbi.1008369.

39. Rafiee S, Kiemel T. Multiple Strategies to Correct Errors in Foot Placement and Control Speed in Human Walking. Experimental Brain Research. 2020-12-01;238(12):2947–2963. doi:10.1007/s00221-020-05949-x.

40. Maus HM, Revzen S, Guckenheimer J, Ludwig C, Reger J, Seyfarth A. Constructing Predictive Models of Human Running. Journal of the Royal Society Interface. 2015;12(103). doi:10.1098/rsif.2014.0899.

41. Nozari P, Finley JM. Development of a Platform to Evaluate Principles of Bipedal Locomotion Using Dynamical Movement Primitives. In: International IEEE/EMBS Conference on Neural Engineering, NER. vol. 2019-March. IEEE Computer Society; 2019-05-16. p. 1062–1065.

